# The interplay between developmental stage and environment underlies the adaptive effect of a natural transposable element insertion

**DOI:** 10.1101/2022.02.09.479730

**Authors:** Miriam Merenciano, Josefa González

**Affiliations:** Institute of Evolutionary Biology (CSIC-Universitat Pompeu Fabra), Barcelona, Spain

**Author notes:** **Correspondence** and requests for materials should be addressed to Josefa González. Josefa González, **Email:**. **Author Contributions:** MM and JG designed research; MM performed research; MM and JG analyzed data; and MM and JG wrote the paper.

**Keywords:** cold-stress, immune-stress, Drosophila, transposable elements

## Abstract

Establishing causal links between adaptive mutations and their ecologically relevant phenotypes is key to understanding the process of adaptation, a central goal in evolutionary biology that is also relevant for conservation biology, medicine and agriculture. Although progress has been made, the number of causal adaptive mutations identified so far is still limited as gene by gene, and gene by environment interactions, among others, complicates linking genetic variation with its fitness–related effects. Transposable elements, often ignored in the quest for the genetic basis of adaptive evolution, are known to be a genome-wide source of regulatory elements across organisms that at times can lead to adaptive phenotypes. In this work, we combine gene expression, *in vivo* reporter assays, CRISPR/Cas9 genome editing, and survival experiments to characterize in detail the molecular and phenotypic consequences of a natural *Drosophila melanogaster* transposable element insertion: the *roo* solo-LTR *FBti0019985*. This transposable element provides an alternative promoter to the transcription factor *Lime*, involved in cold- and immune-stress responses. We found that the effect of *FBti0019985* on *Lime* expression depends on the interplay between the developmental stage and the environmental conditions. We further establish a causal link between the presence of *FBti0019985* and increased survival to cold- and immune-stress. Our results exemplify how several developmental stages and environmental conditions need to be considered to characterize the molecular and functional effects of a genetic variant, and add to the growing body of evidence that transposable elements can induce complex mutations with ecologically relevant effects.

## INTRODUCTION

Establishing causal links between mutations and their relevant fitness-related phenotypes is crucial in biology, with implications for evolution, development and disease (1–3). However, causal genotype-phenotype links are difficult to establish since the effect of a mutation can depend on the genetic background (epistasis) as well as on other contexts such as the environment and the developmental stage (4–6). Because context dependence contributes to diverse traits in diverse organisms, a shift from identifying the impact of a mutation in a particular context to pinpointing the spectrum of effects of a mutation is needed to provide a more realistic picture of the genotype-phenotype map (7).

Identifying adaptive mutations and analyzing how context dependence influences their effects is even more relevant in the current scenario of rapid environmental change (1, 8). To date, most studies that aim at characterizing adaptive mutations have focused on single nucleotide polymorphism (SNP) variants, that are easier to detect by the commonly-used short-read sequencing techniques. However, other types of mutations such as transposable elements (TEs) that are known to be a source of adaptive mutations across organisms, have been understudied so far (9). The increased availability of whole genome sequences and advances in sequencing technologies, such as the improvements in long-read sequencing techniques, are fostering the discovery of candidate adaptive TEs (10). Besides identifying adaptive mutations at the DNA level, and their fitness-related trait in a relevant ecological context, pinpointing the molecular mechanism by which the mutation influences the phenotype is key to conclude that a mutation has an adaptive effect. TEs can affect gene structure and expression through many different molecular mechanisms (11, 12), and some of these changes have been associated with adaptive phenotypes (13–19). In Drosophila, a dual role as enhancer and promoter of an adaptive insertion conferring tolerance to bacterial infection has been recently reported (18). There are also several examples of TE-induced mutations affecting more than one phenotype, such as the disruption of the *Chkov1* gene leading to resistance to pesticides and to viral infection (20, 21). Indeed, *D. melanogaster* is an unrivaled model organism to investigate genotype-phenotype links because it has one of the best functionally annotated genomes and powerful tools to genetically modify the organism *in vivo*, which allows not only to identify the molecular mechanism behind the adaptive effect of a mutation, but also to demonstrate its causality (22).

The *roo* solo-LTR element known as *FBti0019985*, has been previously identified as a candidate adaptive TE insertion: the patterns of nucleotide diversity in its flanking regions suggest that this insertion has increased in frequency due to positive selection (23, 24). *FBti0019985* is inserted in the promoter region of the *Lime* gene, which is located in the first intron of the *cbx* gene (Figure 1). *Lime* is a C_2_H_2_-type zinc finger transcription factor associated with chill-coma resistance and immune response (25, 26), and *cbx* is a ubiquitin-conjugating enzyme that has also been associated with immune-stress (18, 27). Although there is evidence for a role of *FBti0019985* in nearby gene structure and expression, with potential consequences for cold-stress tolerance, these results were based on association studies on a limited number of genetic backgrounds, and the molecular mechanism linking *FBti0019985* and its phenotypic consequences remains unknown (18, 24, 28). Moreover, because *FBti0019985* is inserted nearby genes involved in cold-and immune--stress responses, this mutation could be affecting both phenotypes.

**Figure 1.**
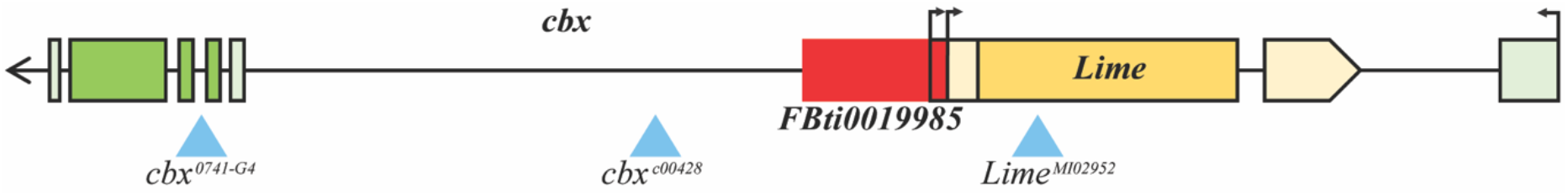
Schematic representation of the genomic region where *FBti0019985* is inserted. *Lime* and *cbx* genes are represented in yellow and green, respectively. Light boxes represent UTR regions and darker boxes represent exons. *FBti0019985* insertion is depicted in red. Transcription start sites, including the one that *FBti0019985* adds, are represented as black arrows. Transgenic insertion sites are represented with blue triangles.

In this work, we characterized in detail the molecular and phenotypic effects of *FBti0019985* under cold-and immune-stress conditions, both of them relevant for the survival of *D. melanogaster* in natural environments. We integrated gene expression analysis, *in vivo* reporter assays, CRISPR/Cas9 genome editing in natural strains, and survival experiments, to establish a causal link between the presence of the insertion and its ecologically relevant phenotypic effects.

## RESULTS

### *Lime* expression changes affect immune-and cold-stress survival and *cbx* expression changes affect immune-stress survival

To provide further evidence for the role of *Lime* in cold-and immune-stress and for the role of *cbx* in immune-stress, we performed survival experiments in these two stress conditions. Previous evidence for a role of *Lime* in immune-stress was obtained in larvae after wasp infection (26). To test whether *Lime* down-regulation is also associated with increased sensitivity to bacterial infection in adult flies, we exposed *Lime* mutant flies to *P. entomophila*. We first confirmed that *Lime^MI02952^* mutant down-regulates *Lime* expression (Fig. 2A, Table S1A and S1B), and we also discarded that *lime^MI02952^* affected the expression of *cbx*, as mutations in this genomic region have previously been suggested to affect expression of both genes (Fig. 2A, Table S1B) (18, 24). As expected, we found that *Lime* mutant flies, both male and females, were more sensitive to *P. entomophila* infection compared with wild-type flies (Fig. 2B, Table S1C). *Lime* is up-regulated in strains selected for cold-tolerance, and up-regulation of *Lime* in embryos is associated with increased egg-to-adult viability in nonstress and cold-stress conditions (24, 25). Consistent with these results, we found that *Lime^MI02952^* flies, which down-regulate *Lime*, showed reduced egg-to-adult viability in nonstress and in cold-stress conditions (Fig. 2C, Table S1D).

**Figure 2.**
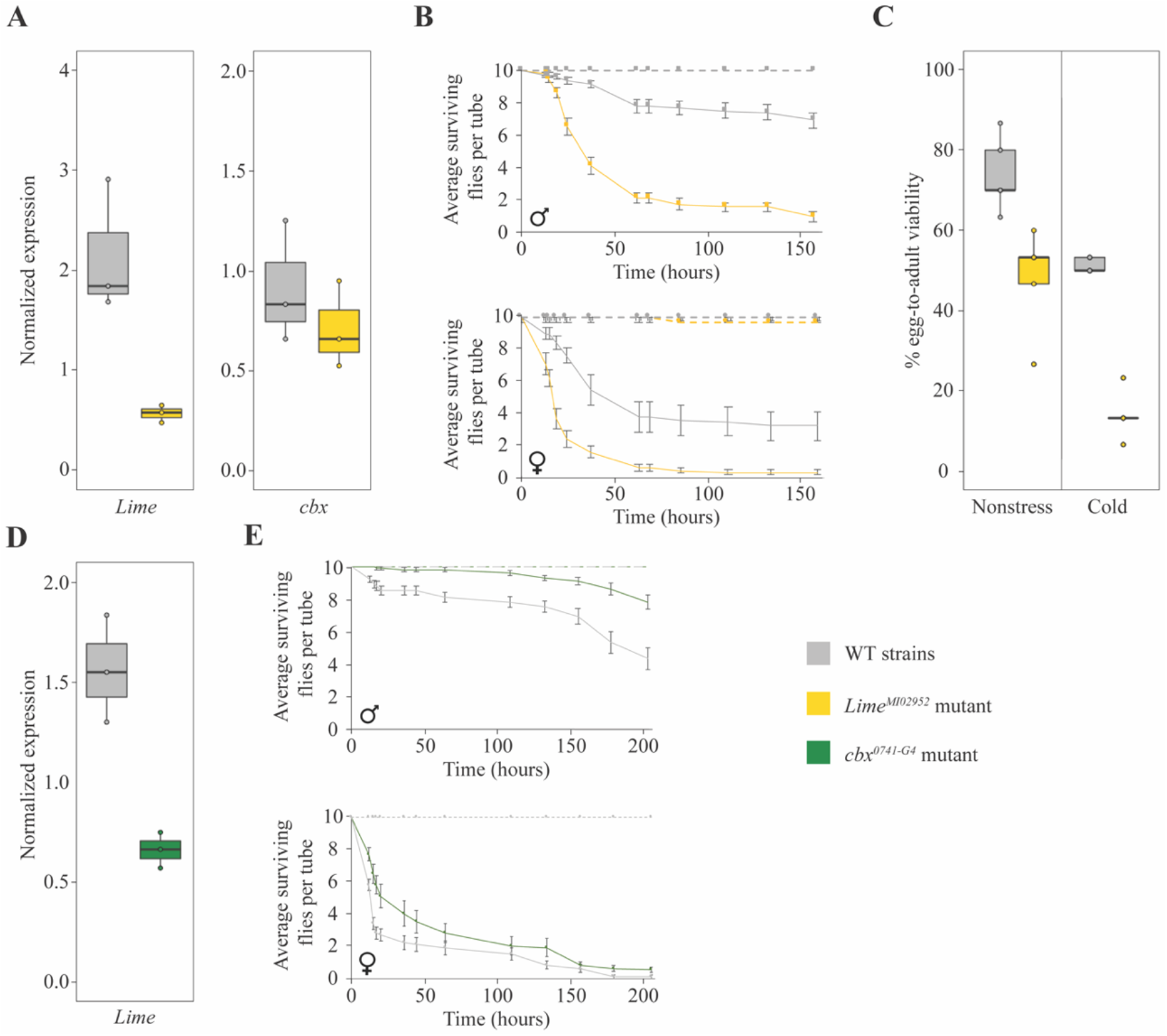
*Lime* mutants are more sensitive to *P. entomophila* infection and have reduced egg-to-adult viability under cold-stress. **A** Normalized expression of *Lime* and *cbx* with *Act5C* in the *Lime^MI02952^* mutant strain compared with the WT strain. **B)** Survival curves of *Lime^MI02952^*mutant males and females compared with the WT strain. Survival curves in non-infected conditions are depicted as dotted lines while survival curves after *P. entomophila* infection are depicted as continuous lines. Error bars represent SEM. **C)** Egg-to-adult viability in nonstress and in cold-stress conditions of *Lime^MI02952^* mutants compared with the WT strain. **D)** Normalized expression of *Lime* with *Act5C* in the *cbx^0741-G4^* mutant strain compared with the WT strain. **E)** Survival curves of *cbx^0741-G4^* mutant males and females compared with the WT strain. Survival curves in non-infected conditions are depicted as dotted lines while survival curves after *P. entomophila* infection are depicted as continuous lines. Error bars represent SEM.

Evidence for a role of *cbx* in immune response is based on the analysis on two mutantstocks: *cbx^c00428^* and *cbx^0741-G4^* (Fig 1). However, although *cbx^c00428^* mutant flies were associated with increased sensitivity to the gram positive bacteria *S. aureus* (27), this mutant does not affect *cbx* expression(Table S1B) (18). On the other hand, we found that *cbx^0741-G4^*, a null mutant associated with increased tolerance to *P. entomophila* infection (18), also down-regulates *Lime* expression (Fig. 2D, Table S1B). These results suggest that *cbx* indeed plays a role in immune response as *Lime* down-regulation alone is associated with increased sensitivity to *P. entomophila* infection (Fig. 2B, Table S1C), while *Lime* downregulation and knockout of *cbx* expression in the same genetic background was associated with increased tolerance to infection (Fig. 2E, Table S1C).

Overall, our results provide further evidence suggesting that changes in *Lime* expression affect cold-and immune-stress responses, and changes in *cbx* expression affects immune-stress response (18, 25–27).

### *FBti0019985* is not associated with changes of expression of *cbx* in response to immune-stress

To test whether *FBti0019985* is associated with changes of expression of *cbx* in response to immune stress, we used two outbred populations, that were fixed for the presence (or absence) of the *FBti0019985* insertion in a heterogenous unlinked background (29, 30). Note that previous evidence for an association between *FBti0019985* and *cbx* expression was based on the analysis of heterozygous inbred strains and concluded that the effect was background dependent (18).

We found that in outbred populations, *cbx* expression was not significantly affected by the insertion genotype (presence/absence of *FBti0019985*), the experimental condition (nonstress *vs*. immune-stress) or the interaction between the genotype and the experimental condition (ANOVA p-values > 0.05; Fig. 3A and Table S2), suggesting that the TE does not affect *cbx* expression. We thus focused on the effect of *FBti0019985* on *Lime* for the rest of this work.

**Figure 3.**
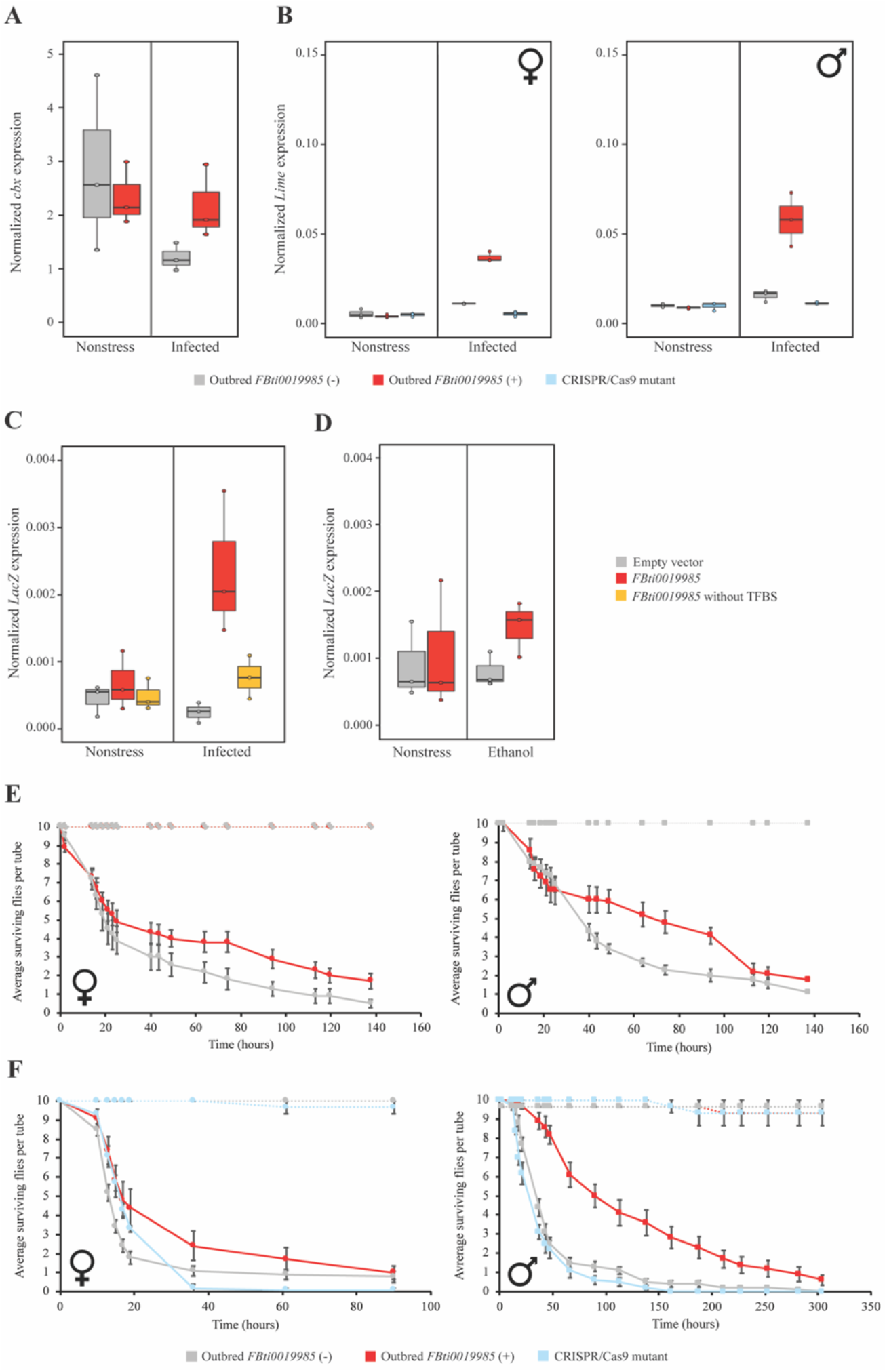
*FBti0019985* is associated with *Lime* up-regulation in guts under immune-stress conditions and with increased tolerance to *P. entomophila* infection. **A)** Normalized expression of *cbx* with *Act5C* in female guts in nonstress and after *P. entomophila* infection. Two-way ANOVA: Genotype, G (presence/absence of *FBti0019985)* p=0.687; Experimental condition, EC (nonstress *vs*. immune stress) p=0.138; Interaction, GxEC p=0.219. **B)** Normalized expression of *Lime* with *Act5C* in nonstress and immune-stress conditions of female and male guts from outbred populations with and without *FBti0019985*, and for CRISPR/Cas9-mutant flies. Two-way ANOVA outbred females: G p<0.001, EC p<0.001, GxEC p<0.001. Two-way ANOVA CRISPR/Cas9 mutant females: G p<0.001, EC p<0.001, GxEC p<0.001. Two-way ANOVA outbred males: G p=0.002, EC p<0.001, GxEC p=0.001. Two-way ANOVA CRISPR/Cas9 mutant males: G p=0.001, EC p<0.001, GxEC p=0.001. **C)** Normalized expression of the reporter gene *lacZ* with *Act5C* in guts under nonstress and immune-stress conditions from transgenic flies. Two-way ANOVA *FBti0019985:* G p=0.009, EC p<0.065, GxEC p=0.026. Two-way ANOVA *FBti0019985* without TFBSs: G p=0.036, EC p<0.024, GxEC p=0.084. **D)** Normalized expression of the reporter gene *lacZ* with *Act5C* in transgenic adult female flies under nonstress conditions and after an ethanol exposure. Two-way ANOVA *FBti0019985:* G p=0.272, EC p<0.669, GxEC p=0.494. **E)** Survival curves of outbred flies with and without *FBti0019985* and **F)** CRISPR/Cas9-mutant flies after *P. entomophila* infection. Survival curves in non-infected conditions are depicted as dotted lines while survival curves after *P. entomophila* infection are depicted as continuous lines. Error bars represent SEM. Log-rank p-values < 0.05.

### *FBti0019985* is the causal mutation inducing *Lime* up-regulation in adult flies under immune-stress

To test for an association between *FBti0019985* and *Lime* expression in nonstress and immune-stress conditions, we used the same two outbred populations described above. While no differences in *Lime* expression levels between outbred flies with and without the insertion were found in nonstress conditions, the outbred population with *FBti0019985* had increased *Lime* expression under immune-stress conditions, both in male and female guts (Fig. 3B, and Table S2). To confirm that the *FBti0019985* is the mutation causing increased *Lime* expression, we measured *Lime* levels in a natural strain with a precise deletion of *FBti0019985* that was generated using the CRISPR/Cas9 technique (see Methods). While no differences in expression levels of *Lime* were found in nonstress conditions, under immune-stress, the CRISPR/Cas9 mutant showed a *Lime* down-regulation compared with the outbred strain with *FBti0019985*, as expected if this insertion is the causal mutation (Fig. 3B and Table S2).

Overall, we found that *FBti0019985* was associated with *Lime* up-regulation under immune-stress conditions, and we established using CRISPR/Cas9 to precisely delete the insertion, that *FBti0019985* was the causal mutation (Fig. 3B).

### *FBti0019985* harbors functional transcription factor binding sites related with immune response

To identify the molecular mechanisms by which *FBti0019985* up-regulates the expression of *Lime* in immune stress conditions, we performed *in vivo* enhancer assays. *FBti0019985* sequence has three predicted binding sites for *DEAF-1*, *tin*, and *Dorsal* transcription factors, which are related with immune response (24, 28). We thus tested whether these immune-related transcription factor binding sites (TFBSs) could be responsible for the *Lime* up-regulation, by using directed mutagenesis to delete them from the *FBti0019985* sequence (see Methods) and cloning this modified *FBti0019985* TE in front of a reporter gene. As expected, the complete sequence of *FBti0019985* drives the expression of the reporter gene only in infected conditions (Fig. 3C, Table S3) (28). When we deleted the three immune-related TFBSs, the expression of the reporter gene was reduced, suggesting that the deleted binding sites were responsible for the enhancer activity of *FBti0019985* in infected conditions (Fig. 3C, Table S3).

To test whether the enhancer effect of *FBti0019985* is stress-specific, we measured the expression of the reporter gene in flies exposed to ethanol-stress. We found no differences in expression under nonstress and ethanol-stress conditions (Fig. 3D, Table S3). These results confirmed that *FBti0019985* is not acting as an enhancer in nonstress conditions in adult flies, and suggested that the enhancer effect of *FBti0019985* is stress-specific.

Besides acting as an enhancer, *FBti0019985* could also be affecting *Lime* expression by adding a new transcription start site (TSS). TEs of the *roo* family, including *FBti0019985*, have previously been shown to add alternative TSS to nearby genes in embryonic stages (31). However, whether the TSS of *FBti0019985* is also used in guts and whether this TSS could contribute to the differential expression of *Lime* in response to immune stress is unknown. We found that the alternative transcript starting in *FBti0019985* was also present in the gut of adult flies both in nonstress and in immune-stress conditions (Table S2). A transcript-specific qRT-PCR showed that the expression level of the transcript that starts in the TE does not increase significantly under infected conditions (t-test p-value > 0.352). Moreover, if we compared the expression level of the transcript that starts in the TE with the total *Lime* expression, we found that its contribution, both in nonstress and immune-stress conditions, is very low: <1% and 4%, respectively (Table S2).

Overall, we demonstrated that *FBti0019985* harbors functional TFBSs related to the immune response that are responsible for its enhancer activity in infected conditions, and that this enhancer activity is stress-specific. Although *FBti0019985* is adding an TSS in the gut, the transcript that starts in the TE does not significantly contribute to the increased expression of *Lime* in infected conditions.

### *FBti0019985* increases tolerance to *P. entomophila* infection

To test whether the up-regulation of *Lime* in outbred flies with *FBti0019985* under immune-stress conditions affects fly survival, we performed infection tolerance assays with *P. entomophila*. In outbred populations, we found that both female and males flies with *FBti0019985* were more tolerant to infection than flies without the insertion (Fig. 3E, Table S4). We repeated the infection tolerance assay including this time the CRISPR/Cas9-mutant strain which has a precise deletion of the *FBti0019985* insertion. As before, we observed that outbred flies with *FBti0019985* were more tolerant to infection compared with outbred flies without the insertion in both females and males (Fig. 3F, Table S4). And, as expected if *FBti0019985* was the causal mutation, the CRISPR/Cas9-mutant strain was more sensitive to infection compared to flies with *FBti0019985* both in females and males (Fig. 3F, Table S4).

### *FBti0019985* drives *Lime* up-regulation in embryos under nonstress conditions

We next tested whether *FBti0019985* affects *Lime* expression in embryos in nonstress and cold-stress conditions. While *FBti0019985* is associated with up-regulation of *Lime* in nonstress conditions, this effect was background dependent, and whether it also affects the expression in cold-stress conditions has not been tested before (24). We confirmed that embryos from the outbred population with *FBti0019985* showed *Lime* up-regulation in nonstress conditions compared to embryos from the outbred population without the insertion (Fig. 4A, Table S2). However, we found no differences in *Lime* expression in cold-stress conditions associated with the presence of the TE (Fig. 4A, Table S2), further suggesting that the effect of the TE is stress-specific. As expected if *FBti0019985* is the causal mutation, CRISPR/Cas9-mutant embryos showed reduced expression levels of *Lime* in nonstress conditions (Fig. 4A, Table S2).

**Figure 4.**
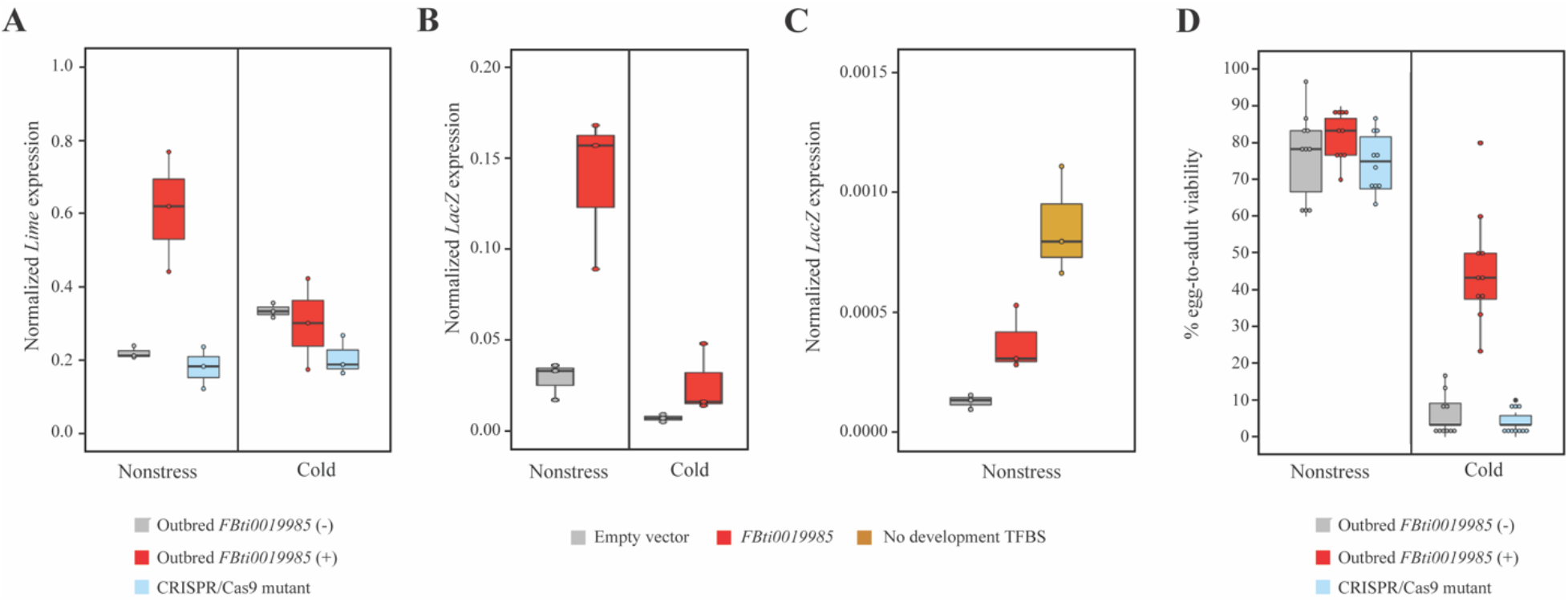
*FBti0019985* is associated with *Lime* up-regulation in embryos and with increased viability in both nonstress and cold-stress. **A)** Normalized expression of *Lime* with *Act5C* in nonstress and cold-stress conditions in embryos from outbred populations, and for the CRISPR/Cas9-mutant strain. Two-way ANOVA outbred: G p=0.018, EC p=0.142, GxEC p=0.007. Two-way ANOVA CRISPR/Cas9 mutant: G p=0.003, EC p=0.056, GxEC p=0.029. **B)** Normalized expression of the reporter gene *lacZ* with *Act5C* in embryos under nonstress and cold-stress conditions from transgenic flies. Two-way ANOVA: G p=0.002, EC p=0.001, GxEC p=0.012. **C)** Normalized expression of the reporter gene *lacZ* with *Act5C* in embryos under cold-stress conditions from transgenic flies. T-test p-value=0.035. **D)** Egg-to-adult viability in nonstress and in cold-stress conditions of outbred populations and CRISPR/Cas9 mutant. Two-way ANOVA outbred: G p<0.001, EC p<0.001, GxEC p<0.001. Two-way ANOVA CRISPR/Cas9 mutant: G p<0.001, EC p<0.001, GxEC p<0.001.

### *Lime* up-regulation in embryos is likely not due to functional binding sites in *FBti0019985* sequence

Consistent with the effect of *FBti0019985* on *Lime* expression in outbred populations, we found that transgenic embryos containing the *FBti0019985* sequence showed significantly increased reporter gene expression only in nonstress conditions (Fig. 4B, Table S3). *In-silico* predictions found TFBSs related with developmental processes in the *FBti0019985* sequence that could be responsible for this enhancer activity of the TE in embryos (28). We thus performed an *in vivo* reporter assay deleting the seven development-related predicted binding sites from the TE sequence (*Dorsal*, *Nub*, *ara/mirr* (x2), *Bap*, *Vnd*, and *Btd*) (see Methods). Contrary to our expectations, we did not find reduced expression of the reporter gene when the binding sites were removed from the *FBti0019985* sequence (Fig. 4C, Table S3). Indeed, we found increase expression of the reporter gene suggesting that, these binding sites might be functional but they were repressing the expression of the reporter gene (Fig. 4C, Table S3).

Finally, the level of expression of the transcript that starts in the *FBti0019985* insertion was not significantly different in nonstress and cold-stress conditions (t-test p-value= 0.174.). In embryos, the contribution to the total *Lime* expression of the transcript that starts in the TE was higher than the one observed in adults (13.6% and 10.2% in nonstress and stress, respectively *vs*. <1% and 4%; Table S2), as expected since *roo* elements are known to provide embryonic promoters (31).

### *FBti0019985* increases egg-to-adult viability under nonstress and cold-stress conditions

We exposed the outbred strains with and without the element to cold-stress conditions and measured egg-to-adult viability. We found that outbred flies with *FBti0019985* had increased egg-to-adult viability under cold-stress (Fig. 4D, Table S5). As expected if *FBti0019985* is the causal mutation, CRISPR/Cas9-mutants showed reduced viability compared with the outbred strain with the element in nonstress and cold-stress conditions (Fig. 4D, Table S5).

## DISCUSSION

In this work, we characterized a naturally occurring TE-induced mutation and found that the effect of the mutation depends on the interplay between the environmental conditions and the developmental stage. While in nonstress conditions, the *FBti0019985* insertion only affects *Lime* expression in embryos, in stress conditions *FBti0019985* affects *Lime* expression in adults, and specifically in response to immune-stress (Fig. 3 and 4). CRISPR/Cas9-mediated deletion of the insertion in a natural population confirmed that the TE is the causal mutation leading to gene upregulation in nonstress in embryos and in immune-stress in adults. Furthermore, we showed that *FBti0019985* is the causal mutation leading to increased bacterial infection tolerance and increased viability in nonstress and cold-stress conditions (Fig. 3 and 4). Previous evidence for the complex effects of TE-induced mutations were mostly associated with coding mutations (32, 33). For instance, *Syncytin-1*, a human-endogenous retrovirus (HERV) gene with a clear role in human placental formation (34, 35), has also been associated with the development of neuropsychological disorders and multiple sclerosis under certain environmental stress conditions such as infection, or drug application (32, 36). Similarly, *Supressyn*, another gene from a retroviral origin also involved in placental development, has been recently linked with resistance to viral infection, demonstrating that its effect is context-dependent (33). Our results thus showed that besides TE-induced coding mutations, the effect of TE-induced regulatory mutations is also complex, and further suggest that TEs could be more likely to have complex effects compared to SNPs (37, 38).

Besides adding an alternative TSS to *Lime, FBti0019985* also acts as an enhancer (24, 31). There is increasing evidence across organisms that regulatory regions can have a dual function as both promoters and enhancers (39, 40). Indeed, in Drosophila, 4.5% of enhancer sequences overlap with TSS (41, 42). TE insertions have been documented to contribute to gene regulation by functioning as enhancers (43, 44) and promoters (31, 45), and recently a dual role as both enhancer and promoter in immune-stress conditions has been described for *FBti0019386*, a natural TE insertion in *D. melanogaster* (18). However, unlike *FBti0019386*, the alternative transcript starting in *FBti0019985* does not seem to have a significant contribution to the changes in gene expression in the stages and conditions analyzed in this work. Thus, characterization of other TEs with the potential to increase transcript diversity and to act as enhancers is needed to evaluate the contribution of this type of mutations to complex regulatory regions.

Our results also suggest that the molecular mechanism by which a TE increases expression of its nearby gene could be different in nonstress *vs*. stress conditions. While we found that the insertion harbors functional binding sites for immune-related transcription factors (Fig. 3C), a mechanism other than adding binding sites appear to be responsible for the enhancer role in embryonic stages (Fig. 4C). Although we cannot discard that binding sites not identified in this work are responsible for the enhancer activity of *FBti0019985* in embryos, other mechanisms such as inducing epigenetic changes could also be responsible for the observed changes in expression (18, 46, 47).

Finally, we speculate that the effect of *Lime* on sugar metabolism might underlay the increased immune-and cold-stress survival observed (Fig 3. and Fig. 4) (26). *Lime* affects the levels of glucose and trehalose, which has been shown to affect immune cell proliferation and activation (26). Moreover, changes in glucose and trehalose have been reported in cold-shock and rapid cold hardening treatments in *D. melanogaster* (48, 49). Thus, increased basal levels of *Lime* expression in embryos might be beneficial when flies are exposed to cold-stress. Indeed, distinct basal transcriptional states have been associated with different phenotypic responses to enteric infection and also to cold-stress in *D. melanogaster* (25, 50). However, further experiments are needed to link the effect of *FBti0019985* on *Lime* expression with changes in sugar metabolism and its subsequent effects on cold and immune tolerance.

Overall, this work provides evidence for *FBti0019985* being the causal mutation responsible for *Lime* up-regulation as a result of the interplay between developmental stage and environment. We further demonstrate that *FBti0019985* increases viability in cold-stress and immune-stress. These results open up the question of how often tissue-specific and environment-dependent gene expression could be due to the presence of TEs and calls for their inclusion in studies aimed at understanding the context-dependent effect of mutations. Furthermore, our results provide support for the need to expand the concept of the genotype-phenotype map from identifying the impact of a mutation in a particular context to exploring the spectrum of effects in different contexts, including different genetic backgrounds, environments, and developmental stages.

## METHODS

### Fly stocks

Fly stocks were reared on standard fly food medium in a 12:12 hour light/dark cycle at 25°C.

### Laboratory mutant and RNAi knock-down strains

We used three laboratory mutant strains and one RNAi knock-down strain that were likely to affect *Lime* and/ or *cbx* genes. We used *Lime^MI02952^* mutants (Bloomington Drosophila Stock Center (BDSC) stock number #36170) that contains a MiMIC insertion in the first exon of *Lime* gene and a RNAi knock-down strain for the same gene (stock number #33735) (Table S1A). We also used *cbx^0741-G4^* mutants (stock number #63767) and *cbx^c00428^* mutants (stock number #10067) with a PiggyBac insertion in the first and third intron of *cbx*, respectively. *Lime* and *cbx* expression levels were measured in all the strains mentioned above and only *Lime^MI02952^* mutants that affected *Lime* expression were used in the phenotypic assays.

### Outbred strains

We used the outbred populations with and without *FBti0019985* generated in Merenciano et al. (2019) (29). Briefly, they were generated by a round-robin cross-design of inbred lines from the Drosophila Genetic Reference Panel (DGRP) (51) and isofemale lines from different European populations (29). Outbred populations were maintained by random mating with a large population size for over five generations before starting the first experiments.

### CRISPR/Cas9 mutant strains

To generate a CRISPR/Cas9 strain with a precise deletion of *FBti0019985* in the natural outbred population we followed a two-step approach. First, we substituted the insertion by the DsRed fluorescence marker and then, in a second step, we removed the visual marker to avoid any possible effect of the introduced DsRed sequence in the results (Figure S1). For the substitution of *FBti0019985* by the DsRed fluorescence marker, guide RNAs (gRNA1 and gRNA2) were designed in the *FBti0019985* flanking region using the flyCRISPR target finder (http://targetfinder.flycrispr.neuro.brown.edu/) and cloned into pCFD5 plasmid following the pCFD5 cloning protocol (www.crisprflydesign.com) (52) using the primers 5’-gcggcccgggttcgattcccggccgatgcaagctagacttatttgagatagttttaga gctagaaatagcaag-3’ and 5’-attttaacttgctatttctagctctaaaaccaga gaaacgtcgagctgcgtgcaccagccgggaatcgaaccc-3’. gRNA1 is 5’-tatctcaaataagtctagct-3’, while gRNA2 is 5’-cagagaaacgtcgagctgcg-3’. A donor DNA containing two homology arms flanking the DsRed sequence for homology repair were cloned into the pHD-ScarlessDsRed plasmid (https://flycrispr.org/scarless-gene-editing/) using the Q5^®^ High-Fidelity DNA Polymerase kit (New England Biolabs). Left homology arm contained the sequence in 2R:9870299-9871095 (FlyBase Release 6) (53) while the right homology arm contained the sequence in 2R:9871529-9872365 (FlyBase Release 6) (53) from the outbred population with *FBti0019985*. To avoid cleavage of the donor construct and mutagenesis after integration by CRISPR/Cas9, two single-nucleotide synonymous substitutions (C > G for sgRNA1; G > T for sgRNA2) were introduced into the two sgRNA target site PAM sequences, respectively. The pCFD5 plasmid containing the gRNAs, the donor pHD-ScarlessDsRed plasmid containing the homology arms, and a plasmid containing Cas9 endonuclease were co-injected as a unique mix into approximately 550 embryos from the outbred population with *FBti0019985*. All the injections were performed using the following mix concentrations: pCFD5 plasmid at 100ng/ul, donor plasmid at 500ng/ul, and Cas9 plasmid at 250ng/ul. Offspring was screened for eye fluorescence. Flies with the desired mutation were backcrossed with the parental line for a minimum of five generations to remove potential off-targets generated during the process (54, 55). Then, an homozygous strain containing the deletion of *FBti0019985* was established. The substitution of *FBti0019985* by DsRed was checked by PCR with two primer pairs: 5’-aacaatgcaagtccgtgctc-3’, 5’-gtggttcctccacccttgtg-3’ and 5’-ggccgcgactctagatcataatc-3’ and 5’-gtggttcctccacccttgtg-3’. PCR bands were confirmed by Sanger sequencing. Then, to remove the DsRed sequence, we applied again the CRISPR/Cas9 technique designing two gRNAs (gRNA3 and gRNA4) in the flanking regions of the DsRed sequence as mentioned before and cloning them into the pCFD5 plasmid using the primers 5’-gcggcccgggttcgattcccggccgatgcttgaacactaatgacaatttgttttagagctagaaatagcaag-3’ and 5’-attttaacttgctatttctagctctaaaacagctcacaactgcgcagctctgcaccagccgggaatcgaaccc-3’. gRNA3 is 5’-ttgaacactaatgacaattt-3’, while gRNA4 is 5’-gagctgcgcagttgtgagct-3’. A donor DNA containing only two homology arms for homology-directed repair were cloned into the pHD-ScarlessDsRed plasmid. Left homology arm contained the sequence in 2R:9870299-9871095 (FlyBase Release 6) (53) while the right homology arm contained the sequence in 2R:9871529-9872365 (FlyBase Release 6) (53) from the outbred population with *FBti0019985* adding two single-nucleotide synonymous substitutions (G > A for sgRNA3 site; C > T for sgRNA4 site). The pCFD5 plasmid containing the gRNAs 3 and 4, the donor pHD-ScarlessDsRed plasmid containing the homology arms, and a plasmid containing Cas9 endonuclease were co-injected as explained before into embryos from the CRISPR/Cas9 mutant strain containing the DsRed fluorescent marker. This time, the offspring was screened for the absence of eye fluorescence. An homozygous strain containing the DsRed deletion was established as mentioned above. The DsRed deletion was checked by PCR using the primer pair 5’-aacaatgcaagtccgtgctc-3’ and 5’-cgtaggatcagtgggtgaaaatg-3’, and finally PCR bands were confirmed by Sanger sequencing. No polymorphism were found in the sequenced region except for the two single-nucleotide synonymous substitutions introduced into the two sgRNA target site PAM sequences.

### Transgenic strains

For the *in vivo* reporter assays, we used the transgenic flies with the *FBti0019985* sequence cloned in front of the *lacZ* reporter gene generated in Ullastres et al. (2021) (18), and we compared them with transgenic strains with the placZ.attB empty vector to control for possible *lacZ* expression driven by the vector sequence itself. Transgenic flies with deleted TFBSs were generated using the placZ.attB vector containing the *FBti0019985* sequence (18), and performing directed mutagenesis for each TFBS with the Q5 Site-Directed Mutagenesis Kit (New England Biolabs). Primers used for deleting the TFBSs can be found in Table S6. The placZ.attB vectors containing the *FBti0019985* and the desired deletions in TFBSs were microinjected at 350-500 ng/ul into a *D. melanogaster* strain with a stable docking site (Bloomington Stock number: #24749). Offspring was screened for red eyes and the insertion of the construct was verified by PCR and Sanger sequencing with the primer pair 5’-ggtgggcataatagtgttgtttat-3’ and 5’-cgacgtgttcactttgcttgt-’3. Three independent homozygous stocks were generated and used as biological replicates for the qRT-PCR experiments.

### Expression analysis

#### Sample collection

##### *P. entomophila* infection

5-7 day-old flies from each strain tested were separated by sex and placed in vials with fresh food in groups of 25-35. We allowed flies to recover from CO_2_ anesthesia for 24 hours at 25°C. To expose the flies to the gram-negative bacteria *P. entomophila* infection, we followed the protocol described in Neyen et al. (2014) (56). Briefly, after two hours of starvation, we transfer 75-105 flies in groups of 25-35 into three vials with fresh food and a filter paper soaked with 120 ul of a solution containing 1.25% sucrose and bacterial preparation adjusted to a final OD600 = 50 for females and OD600 = 150 for males. Flies were kept at 29°C, the optimal temperature condition for *P. entomophila* infection. Simultaneously, a total of 75-105 flies were also transferred in groups of 25-35 into three vials with fresh food and a filter paper soaked with 120 ul of a solution containing sterile LB with 1.25% sucrose and kept at 29°C as a control. Guts from males and from females were dissected after 12 hours. Samples were then flash-frozen in liquid nitrogen and stored at −80°C until sample processing.

##### Cold-stress treatment

5-7 day-old flies from each strain tested were allowed to lay eggs at 25°C in a fly cage with egg-laying medium (2% agar with apple juice and a piece of fresh yeast) for four hours. After these four hours, adults were removed and the plate containing embryos was kept at 1°C for four additional hours. Simultaneously, another plate with embryos was kept at 25°C for four additional hours as a control. 4-8 hour-old embryos were then collected from both plates using the method described in Schou et al. (2013) (57) adding an additional step of dechorionation during 10 minutes with 50% bleach. Finally, samples were flash-frozen in liquid nitrogen and stored at −80°C until sample processing.

##### Ethanol exposure

5-7 days-old flies from each strain tested were separated by sex and placed in vials with fresh food in groups of 25. We allowed flies to recover from CO_2_ anesthesia for 24 hours at 25°C. To expose the flies to ethanol we followed the method described in Maples et al. (2011) (58). Briefly, flies were transferred to an empty vial and a coated cotton ball with 0.5 ml ethanol was inserted into the exposure vial for 9 minutes ensuring that the alcohol was facing into the vial and not towards the wall. Simultaneously, coated cotton balls with 0.5 ml H_2_O were inserted into the control vials. After that, flies were flash-frozen in liquid nitrogen and stored at − 80°C until sample processing.

### RNA extraction and cDNA synthesis

RNA was extracted using the GeneEluteTM Mammalian Total RNA Miniprep Kit following manufacturer’s instructions (Sigma). RNA was then treated with DNase I (Thermo). cDNA was synthesized from a total of 250-1,000 ng of RNA using the NZY Fisrt-Strand cDNA synthesis kit (NZYTech).

### qRT-PCR analysis

*Lime* expression was measured using the forward primer 5’-gagcagttggaatcgggttttac-3’ and the reverse primer 5’-gtatgaatcgcagtccagccata-3’ spanning 99 bp cDNA in the exon 1/exon 2 junction. *cbx* expression was measured with the forward primer 5’-gggaaaacgatctgggagca-3’ and the reverse primer 5’-gtcggagaagttgagtggga-3’ spanning 233 bp cDNA in the exon 2/exon 3 junction. *lacZ* reporter gene expression was measured using the forward primer 5’-cctgctgatgaagcagaacaact-3’ and the reverse primer 5’-gctacggcctgtatgtggtg-3’. Gene expression was normalized with *Act5C* (5’-gcgcccttactctttcacca-3’ and 5’-atgtcacggacgatttcacg-3’ primers). We performed the qRT-PCR analysis with SYBR Green (BioRad) or with the qPCRBIO SyGreen Mix Lo-Rox (PCRBiosystems) on iQ5 and CFX384 Thermal cyclers, respectively. Results were analyzed using the dCT method (59).

### Transcript start site detection

We performed RT-PCRs to detect whether *FBti0019985* is adding an alternative TSS to the *Lime* gene in outbred flies carrying *FBti0019985* after different stress conditions. We used the forward primer 5’-aaaactcaacgagtaaagtcttc-3’ and the reverse primer 5’-tataaagttccaacgcccagc-3’ to detect the *Lime* transcript starting in the TE. The forward primer 5’-cgcagagaaacgtcgagctg-3’ and the reverse primer 5’-cacgttaaattcactagggtggc-3’ were used to detect *Lime* total transcript. Outbred population without *FBti0019985* was used as a control sample.

### Phenotypic assays

#### *P. entomophila* infection

100 5-7 day-old male flies and 100 5-7 day-old females from each one of the strains tested were infected with the gram-negative bacteria *P. entomophila* as described before. Simultaneously, a total of 30 male and female flies were tested as controls. We counted the number of dead flies in every vial at different time points for at least 6 days. Log-rank tests were performed to analyze survival curves with SPSS v21 software.

#### Egg-to-adult viability under cold-stress

5-7 day-old flies from each strain tested were allowed to lay eggs for 4 h at 25°C in a fly cage with egg-laying medium (2% agar with apple juice and a piece of fresh yeast) for four hours. Then, adults were removed from the cage and plates were kept for four additional hours at 25°C. After that, 4-8 hour-old embryos were collected using the method described in Schou et al.

(2013) (57) and placed in vials with fresh food in groups of 30. In total, 240-300 embryos were tested for each strain. Cold-stressed vials were kept at 1°C for 15 h and then maintained at 25°C until adult emergence. Simultaneously, control vials were kept at 25°C and never exposed to cold. Percentage egg-to-adult viability was calculated based on the number of emerged flies to the total number of embryos placed in each vial. Statistical significance was calculated performing ANOVA using SPSS v.21 combining all the data into a full model: experimental condition (stress and nonstress), insertion genotype (presence/absence of the insertion) and interaction between these two factors.

## DATA AVAILABILITY

The data is available within the Article, Supplementary Information or upon request.

## ACKNOWLEDGEMENTS

We thank Fillip Port from the Division of Signaling and Functional Genomic led by Prof. Dr. Michael Boutros for his advice in the generation of the CRISPR/Cas9 mutants. This project has received funding from the European Research Council (ERC) under the European Union’s Horizon 2020 research and innovation programme (H2020-ERC-2014-CoG-647900).

